# Genome-Wide Association Studies of malaria susceptibility and resistance: progress, pitfalls and prospects

**DOI:** 10.1101/456707

**Authors:** Delesa Damena, Awany Denis, Lemu Golassa, Emile R. Chimusa

## Abstract

*P. falciparum* malaria is still among the leading causes of child mortality in sub-Saharan Africa; killing hundreds of thousands of children each year. Malaria has been recognized as one of the prominent evolutionary selective forces of human genome that led to the emergence of multiple host protective polymorphisms associated with minimizing the risk of developing severe malaria in endemic areas. A comprehensive understanding of the genetic bases of malaria resistance can shed light to the molecular mechanisms of host-parasite interactions that can potentially pave ways to the development of new therapeutics and vaccines. Genome-wide association studies (GWASs) have recently been implemented in malaria endemic areas and identified a number of novel association genetic variants. Despite this success, only few variants did replicate across the studies and the underlying biology is yet to be understood for the majority of the novel variants. Besides, there are several open questions around heritability, polygenic effects, epistatic interactions, genetic correlations and associated molecular pathways among others. In this review, we first assess the progress and pitfalls of malaria susceptibility GWASs. We then, provide an overview of the current progress in post-GWAS approaches and discuss how these approaches can potentially be implemented in malaria susceptibility GWASs to extract further functional information. We conclude by highlighting the importance of multi-step and multidimensional integrative studies for unravelling the genetic basis of malaria susceptibility and resistance at systems biology level.

## Introduction

*Plasmodium falciparum*, the causative agent of the severe malaria, has been infecting humans for at least 5000-10,000 years following the advent and expansions of agriculture ^1–3^. Malaria still poses a huge social, economic and health problems in several low-income countries, particularly in sub Saharan Africa^4,5^. *P. falciparum* infects millions and kills hundreds of thousands of African children each year. However, this constitutes only small proportion (1%) of the populations in endemic areas in which the infections progress to severe malaria such as profound anemia or cerebral malaria^6,7^. The genetic basis of such variations in malaria disease severity in endemic areas is mainly unclear as the well-known Mendelian variants only explain 2.5% of heritability of malaria protections^8^.

Comprehensive understanding of the genetic basis of resistance and susceptibility to severe malaria is crucial to understand molecular mechanisms of host-parasite interactions that can inform the development of effective therapeutics, vaccination, diagnostics and risk prediction strategies^8,9^. In a broader context, the findings from malaria susceptibility studies may also provide new insight to human physiology and genetic susceptibility to other common diseases because ‘malaria is one of the strongest evolutionary forces that shape human genome polymorphisms^9^.

To better understand the underlying genetics, GWASs have recently been implemented in malaria endemic areas and replicated the well-known variants including *HbS* and *ABO* blood group ^10–13^. Despite this success, only few novel variants were identified; suggesting that some of the true association signals might have been attenuated. Which may be resulted from several confounding factors such as: the high genetic diversity of population in malaria-endemic areas, differences in frequencies and effect sizes of protective alleles across populations and the inherent limitations of GWAS approaches among others.

One interesting observation in these studies is the identification of several association signals distributed across the genome that didn’t pass GWAS significance threshold; suggesting the existence of polygenic effects ^10–13^. This raises several key questions including 1) What is the genetic architecture of malaria protection? 2) What is the contribution of polygenic effects? 2) What is the proportion of heritability explained by cell-types, chromosomes, functional groups and molecular pathways? 3) What is the extent and pattern of epistasis and pleiotropy at genome wide scale?

Here we review the current status of malaria susceptibility GWASs and provide guidance to future research directions. We begin by assessing the progress, pitfalls and opportunity of malaria susceptibility GWASs. We then provide an overview of the recent progresses in post-GWAS approaches and discuss how these methods can be implemented in Malaria susceptibility studies to better understand the underlying biology. We conclude by discussing on research areas where further works are needed in light of the global malaria eradication efforts.

## Malaria susceptibility GWASs: progress

The GWAS approach is continuing to provide unprecedented successes in genetic studies of wide ranges of complex-traits and common diseases^14,15,16^. In malaria susceptibility studies, GWAS approach was initially planned to address the acute limitations of the conventional studies, approaches such as candidate gene and linkage analysis that have been implemented in malaria research for more than two decades ^1,17^. The candidate gene-based studies ^^18–20^ and the family based linkage studies 21,22^ have identified several genomic loci associated with malaria susceptibility/resistance. However, the majority of the findings were discordant and failed to be replicated in different populations; mainly because of the their limited power^1^. Thus, in view of understanding the genetic basis of malaria susceptibility and resistance at genome wide scale, a global partnership of malaria researchers, named as Malaria Genomic Epidemiology Network (MalariaGEN)^17^, has been established and successfully conducted multi-center-scale GWASs ^10–13^.

The GWASs have replicated the well-known variants including *HbS* and *ABO* blood groups which reinforce the importance of erythrocyte variants for protection against severe malaria. Besides, the GWASs have identified novel association variants (Table 1) that may directly or indirectly influence the disease outcome. The epidemiology and biology of the well-known variants were reviewed else-where^1^. Below we characterize the novel malaria susceptibility genetic variants identified by GWASs. We first discuss the biology of two variants such as *ATP2B4* and cluster of the *glycophorin genes (GYPA/B/E)* that were convincingly associated and independently replicated across malaria endemic populations and extend our discussion to other potential association signals which didn’t replicate and/or did not reach the GWAS significance threshold.

**Table 1.**
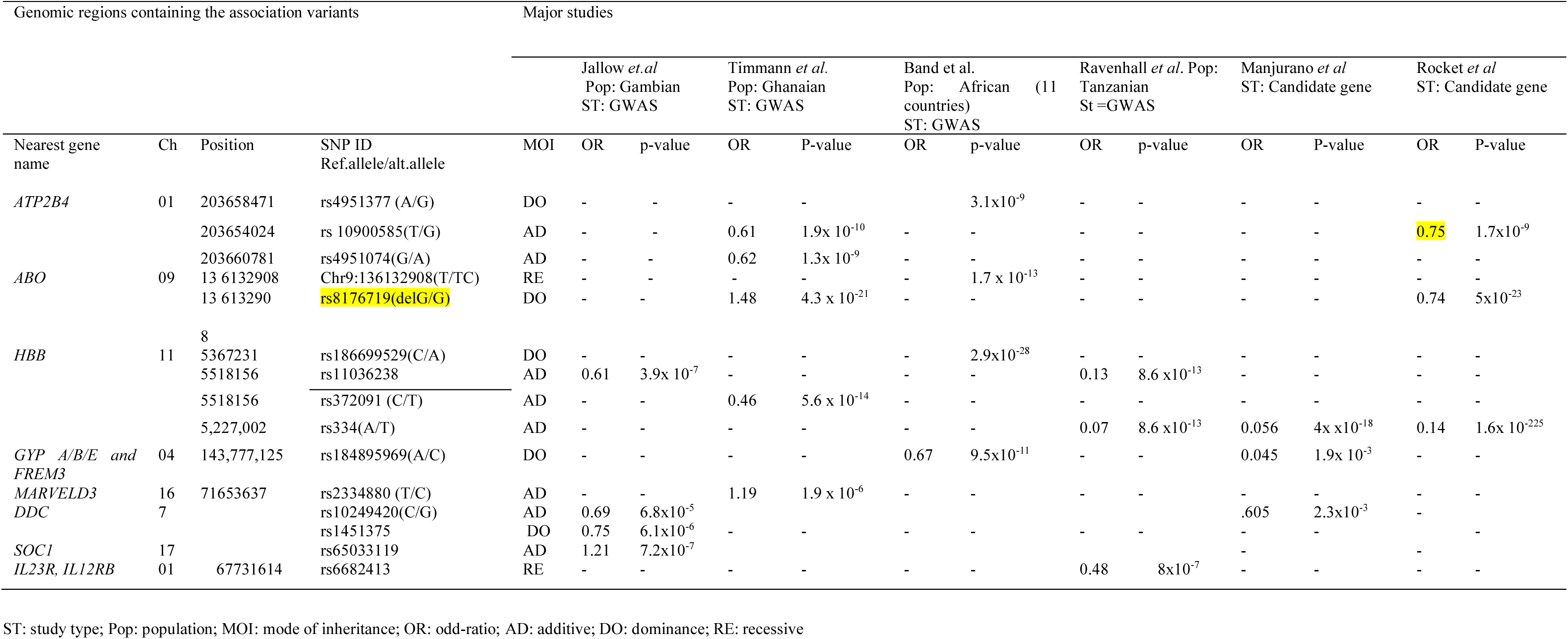
Summary of genetic variants associated with resistance and susceptibility to severe Malaria identified by GWASs

## Novel association variants

### ATP2B4

The association of *ATP2B4* gene with severe malaria was first identified by Timmann *et al*. ^11^in Ghanaian populations. In this study, several SNPs within the *ATP2B4* gene, located on chromosome 1q32 showed significant associations. The protective role of the variants of *ATP2B4* gene was subsequently reported in other studies^12,23^. *ATP2B4* encodes a ubiquitous plasma membrane calcium-transporting *ATPase4* (*PMCA4b*)^24^. *PMCA4b* is widely expressed in the majority of human tissues and is the main transporter of *Ca*^2+^ in erythrocyte membrane^24^. A recent study showed that individuals who have a previously unrecognized haplotype named as ‘haplotype 1’ in their *ATP2B4* gene exhibit a reduced *PMCA4b* expression level^25^. Several SNPs implicated in malaria protection are localized in this haplotype. Timmann *et al*^11^. proposed that the disturbance of intracellular *Ca*^2+^ homeostasis caused by impaired *PMCA4b* regulation might play an important role in interfering with pathological effects of malaria in different tissues. The disturbance of intracellular *Ca*^2+^ homeostasis in RBCs might impair the parasite development and reproduction in a parasitophorous vacuole membrane (PVM)^26^. The dis-regulation of intracellular *Ca*^2+^ in other tissues such as placenta, platelets and brain might modify the adhesion phenotypes and hence minimize adhesion-associated pathologies^27^.

Protective mechanisms of *ATP2B4* variants might also be associated with impaired regulations of nitric oxide (NO), one of the important molecules in malaria pathogenesis and protections ^28^. Increased level of NO was implicated in protection against cerebral malaria while it increases the risk of uncomplicated malaria^29^. In neuronal cells, *PMCA4* modulates the activity of neuronal nitric oxide synthesis (nNOS) which in turn regulates NO bioavailability in calcium dependent fashion^30^. Thus, the reduced production of *PMCA4* might affect the activities of the *nitric oxide synthesis* enzyme in the brain which might affect the concentration of NO and hence protection against severe malaria.

### Cluster of the 3 glycophorin genes (GYPA/B/E)

The largest multi-center malaria susceptibility GWAS which included eleven populations was conducted by Band *et al.*^12^. In this study, 34 genomic regions containing potential susceptibility loci for severe malaria were identified. Among which, a strong signal was observed at locus between *FREM3* gene and cluster of 3 *glycophorin genes (GYPA/B/E*) on Chromosome 4. A haplotype (at SNP *rs184895969)* within this region was reported to reduce the risk of developing severe malaria by about 40% and is common in Kenyan populations with allele frequency of about 10% ^12^.

In addition to the signals near glycophorin genes, six other putative loci that play key roles in membrane biology were identified in this study. A subsequent study in the same populations identified a large number of copy number variants which are characterized by deletion, duplications and hybrid structures in *GYPA* and *GYPB* genes ^31^. Of which a distinct variant called *DUP4* was reported to reduce the risk of severe malaria by about 40% in eastern African (Kenya) populations. Further characterization showed that, this variant is composed of complex *GYPB-A* hybrid and encode *Dantu* antigen in MNS blood group system ^31^. The association of this region with severe malaria was supported by another recent study in Tanzanian populations^32^. The glycophorin gene cluster, *GYPA* and *GYPB* encode the MNS blood group system and are known to be receptors for *P. falciparum* during RBC invasion^33^. *GYPA* and *GYPB* serve as an erythrocyte membrane receptor for *EBA-175* and *EBL-1* proteins of the parasite respectively^34^. This genomic region is also known to be under an ancient selective pressure resulted from host-pathogen arm races between *P. falciparum* and humans^35^. Further functional analysis is required to better understand how these variants affect the invasion and/or development of the parasites in erythrocytes and convey protection against severe malaria.

## Notable novel association variants

### SCO1 and DDC

Notable association signals were identified by the first malaria susceptibility GWAS conducted in Gambian population^10^. The first lead SNP (*rs6503319*) is located close to *SCO1 (synthesis of cytochrome c oxidase)* gene on chromosome 17p13. *SCO1* is a multi-functional signaling protein which plays an essential role in *mitochondrial cytochrome c oxidase (COX)* copper delivery pathways^36^. *COX* catalyzes electron transfer from reduced cytochrome c to oxygen and is abundantly expressed in muscles, brain and liver ^36^. Deficiency of *COX* caused by mutations in *SCO1* gene can lead to respiratory distress and severe metabolic acidosis^37^ which are also the major complications during cerebral malaria ^38^. Further studies are needed to understand how the variants in *SCO1* gene are associated with the pathological pathways of cerebral malaria.

The second notable association signal identified in this study was *Dihydroxypheny-alanine decarboxylase (DDC)* on Chromosome 7p12.2. A recent study in Tanzanian populations replicated the association of *DDC* variants with cerebral malaria ^32^. *DDC* gene encodes *Aromatic-L-amino-acid decarboxlase* enzyme which is involved in biosynthesis of neurotransmitters such as dopamine and serotonin^39^. *DDC* is an essential enzyme for brain and nervous developments and its deficiency is associated with reduced cognitive functions ^39^. *DDC* is involved in cellular immunity and contributes in protection against parasitic disease in invertebrates^40^. Furthermore, mutations in *DDC* gene was reported to be associated with refractoriness of *Anopheles gambiae* mosquito against *P. falciparum* parasites^41^

### MARVELD3

In addition to the *ATP2B4*, Timmann *et al.* ^11^ identified an association SNP (*rs2334880*) on chromosome 16p 22.2 which is linked to *MARVELD3*. However, this association has not been replicated in other studies. *MARVELD3* is one of the components of tight junction proteins in several epithelial and endothelial tissues and is expressed as two alternative spliced variants^42^. These proteins are involved in assembly, development, maintenances and regulations of tight junction. Tight junctions play a major role in intracellular adhesions and involved in sub-cellular signaling mechanisms^42^.

### IL-12 receptors and IL-23 receptors

The most recent malaria susceptibility GWAS was conducted in Tanzanian population^13^. In this study, notable associations signals were identified in immune pathways including in interleukin receptors (*IL-23R and IL-12RBR2*), *in ketch-like proteins (KLHL3)* and Human Leucocyte Antigen (HLA) regions. *Interlekuen-12* is formed from a hetrodimer of *IL12B (ILp40 subunit)* and *IL-12A (ILp35 subunit*)^43^. *IL-12* plays a vital role in stimulating cell-mediated immune response against intra-cellular pathogens through binding to high affinity *IL-12RB1* and *IL-12RBR2* receptor complexes. It promotes the development of *T-helper cells* (*Th1*) and enhances the production of *INF-γ*, which are known to mediate the clearance of intracellular pathogens^43^. In malaria, *IL-12* has been implicated in mediating the protective immunity both in experimental animals and humans^44^. *IL-23* is an important pro-inflammatory cytokine that shares p40 subunits with *IL12*^45^. It induces the differentiation of naive *CD4 T*-cells to *IL-17* which plays key roles in pathogenesis of autoimmune diseases^46^. HLA is encoded by the Major Histocompatibility Complex (MHC), the most polymorphic genes known in human genome. The diversity of MHC is believed to be driven by selection pressure from infectious pathogens and known to be associated with the risk of several infectious diseases^47^. HLA variants such as HLA class I antigen (*HLA-Bw53*) and HLA class II variant (*DRB1*1302-DQB1*0501*) were reported to confer protections against severe malaria in Gambian populations^47^. HLA class I antigen is expressed by liver cells suggesting that T cells (CTL) responses might efficiently act against the liver stage of malaria parasite in individuals with *HLA-Bw53*^47^. Similarly, individuals with *DRB1*1302- DQB1*0501* variant might possess efficient antigen presentation mechanism that can lead to rapid clearance of blood stage parasite^47^.

Variants in immune pathways are of great interest because of their potential to inform the development of effective malaria vaccines ^1^. The current study is interesting in that several putative variants in immune pathways were identified. However, the power of this study is limited because of relatively smaller sample size and weak significance threshold used to interpret the findings. Therefore, further studies with higher detective power are needed to consolidate the findings.

## Challenges of malaria susceptibility GWASs

Despite its promising progress, GWAS approach has several limitations including 1) Weak performances in genetically diverse populations^8^, lack of translation of associated loci into suitable biological hypotheses^16^, 3) the well-known problem of missing heritability^15^, 4) the lack of understanding of how multiple modestly associated loci within genes interact to influence a phenotype^48,^ 5) inefficiency in distinguishing between inflation from bias (cryptic relatedness and population stratification) and true signal from polygenicity^49^, 6) the imperfection of asymptotic distribution of current mixed model association or logistic regression in the specific case of low-frequency variants ^50^. The discussion on limitations of GWAS approaches is beyond the scope of this review. Here we focus on the major challenges of malaria susceptibility GWASs and highlight recent positive progresses.

### Dealing with genetic diversity of African populations

There is high level of genetic diversity and weak linkage disequilibrium (LD) in Africans compared to non-African populations^51,52^ which is believed to be due to adaptation to diverse climate, diet and response to infectious diseases such as malaria and tuberculosis^51,53^. These distinct genetic characteristics created major setbacks to GWASs in African populations ^8^. The fundamental rationale of the GWAS is a ‘common disease common variant’ hypothesis which predicts that common diseases are influenced by common genetic variants that are prevalent in higher frequencies in the populations ^16^. To capture optimum amount of the common variations in human genome, commercial arrays that constitute representative SNPs (tag SNPs) are utilized. The selection of tag SNPs (subset of common SNPs) for genotyping chips is based on LD information derived from reference panel in which African populations are not fairly represented^54^.

Consequently, the African GWASs might have missed several true association signals. For instance, in the first malaria GWAS^10^, *HbS* locus, a well-known variant conferring resistance to severe malaria demonstrated a weak signal (p-values ~10^-7^) because of the weak LD between causal variants and the SNPs that were genotyped. After which authors performed targeted sequencing of the locus, undertook multipoint imputation, used proper reference panel and dramatically improved the signal to p-value ~10^-14^. It was estimated that a GWAS of 0.6 million SNPs based on HapMap phase 1 dataset in European population has an equivalent power to the chips with 1.5 million SNPs in African populations ^54^ However, the coverage of commercial genotyping arrays has been enormously improved to be able to capture the genetic diversities among global populations following the recent technological advances and availability of diverse reference data sets. For instance, Omni microarrays based GWASs were shown to be powerful in studying African populations ^52^. Considerable progresses have been made in bringing African population genomes to the public domain. For instance, different ethnic groups from countries such as: Nigeria, Kenya, Gambia, Ghana and Malawi were included in the International Hapmap 3 Consortium^55^ and 1000 Genomics project consortium that provided a sufficiently deeper sequence for all the major human ancestries ^56^. A more diverse set of African population that includes genomic data of 18 ethno-linguistic groups was recently published by The African Genome Variation Project (AGVP) ^52^. The Consortium on Asthma among African ancestry Population in America (CAAPA)^57^ is also an additional resource that can facilitate genomic analysis in populations of African ancestry. The improvement in reference panels have facilitated imputation-based studies in African populations. For instance, Band *et al.*^9^ showed the feasibility of multi-point imputation based meta-analysis in for malaria GWASs using HapMap3 haplotype panel. However, imputation accuracy was relatively lower compared to that of European populations. Another study showed a substantial improvement of imputation accuracy by using the more diversified AGVP WGS reference panel^52^. The reference dataset is expected to grow further and accelerate genomic research by including wide range of haplotype diversity in African populations.

### Sample size

In GWAS, a stringent p-value (1x10^-8^) is usually needed to declare evidences of genuine associations to minimize false discovery rate that can arise from multiple testing.^58^ Thus, very large sample size is required to achieve genome-wide significance threshold particularly for loci with modest effect sizes. The required sample size is even much higher for studies in population of African ancestry because of the higher genetic diversity. However, the sample sizes of GWASs in African populations are generally small compared to non-Africans ^59^.

In malaria susceptibility GWASs, the success rate of identifying significant associations seems to be a function of sample sizes. in the first Malaria GWAS^10^ that involved 2500 children, none of the association signals passed the GWAS significance threshold; whereas in the subsequent GWASs that involved relatively larger sample sizes ^11,12^, novel association variants were convincingly identified. To this effect, MalariaGen consortium has increased the sample size and included more study populations in its recent GWASs^12^. However, one of the challenges of the MalariaGEN dataset is that the sample sizes obtained from each study population is limited; making it difficult to undertake separate GWASs for specific geographic areas. Allelic heterogeneity of the malaria protective variants is very common in endemic areas. For instance, *HbC,* one of the malaria protective alleles, is common in some parts of west Africa such as Burkina Faso, Ghana, Togo and Benin while absent in other west African countries such as Cameroon and Chad^60^. The sickle cell allele, *HbS*, is known to have different haplotype structure and effect sizes in different regions across sub-Saharan Africa^52^. A recently identified glycophorin variants are common in east Africa and absent in some populations in west Africa^12^. This may suggest that the majority of variants driving the malaria clinical severity are heterogeneous; pressing the need of increasing sample sizes for each study populations.

### Genetic architecture of malaria susceptibility/resistance

The performance of GWASs is dependent on the genetic architecture of the disease/trait under investigation. For the majority of complex diseases/traits, the GWAS variants identified thus far, only explain a very small proportion of heritability; a phenomenon commonly termed as ‘missing’ heritability^15^. There have been different explanations for the ‘missing’ heritability including common disease rare variant hypothesis^61,^ none-additive components, primary epistasis ^62,63^ and polygenic genetic architecture^16^.

One of the challenges of malaria GWASs is that we don’t know much about the genetic architecture of malaria protection trait. First, as one of the prominent evolutionary selective forces, the majority of malaria protective alleles might have evolved under positive selection and might potentially be balanced by other forces^64^. In this case, the protective variants are expected to have large effect sizes and higher allele frequencies which can be detected by the conventional GWAS approaches; provided that proper reference panel and genotyping array are used^^64^. Second, similar to the genetic architecture of other infectious diseases 65^, malaria protection trait might largely be attributed to few rare variants of large effect sizes. In this, the GWASs are underpowered as rare variants might not be in LD with common variants and can’t not be ‘tagged’ by SNPs chip. Third, malaria protection trait might be mainly under pyogenic and epistatic control as suggested be several by authors^1,7,66^ which the conventional GWAS approach can’t capture. Here, we focus on polygenic genetic architecture and epistasis which have been implicated for the ‘missing’ heritability of malaria susceptibility GWASs.

## Polygenic architecture and epistasis: Presenting the absent in the current malaria GWASs

### Polygenic genetic architecture

Polygenic view of genetic architecture is gaining ground in genetic epidemiological studies and widely implicated for the ‘missing’ of heritability in GWAS analysis^67^. The rationale behind polygenic inheritance is that complex-traits/diseases are influenced by multiple variants with modest effects that are too small to pass the stringent genome wide significance threshold^68^. In standard GWAS analysis, ‘Genomic control’ (GC) method is applied as a quality-control measure to minimize spurious associations that can be caused by population structures such as population stratification and cryptic relatedness.

However, a slight inflation of the test statistics which cannot be corrected by GC was initially observed across the genome in a Schizophrenia GWAS^69^. Subsequently, this observation has been supported by other studies^70^and led to the development of a number of statistical tools aiming to capture polygenic signals at genome-wide scale including 1) polygenic scoring method implemented in PLINK software^71^ ; 2) Mixed Linear Models (MLM) such as: GCTA^70^, BOLT-LMM^72^, Bayes R^73^ and LDAK^74,75^, PCGC^76^ 3) linkage-disequilibrium (LD) score regression method^77^ among others.

In polygenic scoring method, two GWAS sample sets are used. First, a ‘training sample’ data set is used to estimate the effect size of each SNPs. Then, subset of SNPs with top ranking p-values are selected to calculate polygenic score in an independent dataset called ‘target sample’ for each individual based on the weighted sum of risk alleles at selected SNPs. The detail of this approach is explained in Purcell *et al.*^69^. The MLM approaches, that jointly accounts for fixed and random effects, is becoming a standard method for estimating heritability of complex diseases/traits. Yang et al.^70^ introduced an elegant method that builds genetic relationships matrix (GRM) of unrelated individuals by simultaneously fitting all SNPs and estimates the proportion of phenotypic variations explained by genotypic variation using restricted maximum likelihood (REML) method. Variations of MLM approaches with different flavors have subsequently been published and implemented for estimating and stratifying heritability of complex traits/diseases in to chromosomes, functional groups and pathways^74–76^. The use of unrelated individuals has an advantage of minimizing the bias from shared environments in pedigree studies.

Another approach is LD score regression method^77^. The rationale of this approach is that there are three sources of variations in GWAS estimates such as true signals, population structure and estimation errors. The LD score regression effectively decomposes these components by exploiting the LD strength of a given SNP with other SNPs. The marginal association is higher for SNPs that are in LD with several variants. The detailed description of the polygenic analysis methods is reviewed in^67^.

### Polygenic contributions in malaria susceptibility/resistance

Co-evolution of host-pathogen model predicts that multiple host loci are involved in resistance/susceptibility to infectious diseases due to the complex interactions between the multi-locus parasite genotype and the corresponding defense from the host-genome ^63,78^. Indeed, malaria might have left multiple genetic variants; the majority of which have effects too small to be detected by the standard GWASs. The existence of polygenic inheritance in malaria protection was predicted by several authors^1, 7^ and supported by GWASs. For instance, the largest malaria susceptibility GWAS so far, identified 34 regions of the genome containing variants with evidence of associations^12^. Earlier GWAS in Ghanaian population identified 40 genomic regions containing 102 SNPs with evidences of association in the discovery phase of the study^11^. The recent GWAS in Tanzanian populations^13^ identified 2322 SNPs at several genomic regions by relaxing the significance threshold to 1x 10 ^-5^. Thus, implementation of polygenic analytic methods in malaria studies may potentially shed more light to the underlying biology. For instance, heritability can be estimated and partitioned in to different cell-types and functional groups and molecular pathways which enable to localize causal variants. Furthermore, these approaches can be extended to explore genetic correlation between susceptibility to malaria and to other infectious diseases. The existence of shared genetic basis between infectious diseases susceptibility/protection is well-documented^, 79^. However, the extent and pattern of genetic correlation has not been systematically investigated at genome-wide scale; partly because of inadequate GWAS data for infectious diseases. Such studies can potentially provide clues to common molecular processes between resistance/susceptibility to infectious diseases that will have practical importance such as: designing multi-purpose vaccine and genetic risk prediction strategies.

### Heritability of severe malaria in Gambian population

To figure out how polygenic analysis can be implemented in malaria susceptibility, we accessed the Gambian malaria susceptibility GWAS dataset from European Phenome Genome Archive (EGA) through data application procedure and estimated heritability of malaria susceptibility using MLM approaches. The Gambian GWAS data is the largest MalariaGen dataset obtained from a single country comprised of 4920 samples (2429 cases and 2491controls) and 1.6 million SNPs that passed GWAS quality control (QC). We first excluded the known malaria susceptibility associated loci and performed stepwise extra QC filtering. Specifically, we focused on sample relatedness, SNP missingness proportion and SNP differential missing proportion which are well-known to affect the accuracy of heritability estimation^75^. We then estimated the heritability using GCTA model for different QC thresholds by including 10 principal components (PCs) as fixed effects to account for population structure. As expected, the estimation is highly unstable when less stringent QC thresholds is applied (varying from 37.8% to 20.1%) as shown in Table 2. However, when more stringent QC (Relatedness threshold (5%), SNP differential missingness proportion (p ≤ 1x 10 ^-3^) and SNPs missing proportion of (p>0.02)) is applied, the estimation became stable (~20.1%, SE =.07). Neither the inclusion of more PCs (15, 20) as fixed effects nor SNP phasing further brought down the estimate. Using the same stringent QC threshold, the estimation was approximately the same for Mandinka ethnic group (~24.3%, SE=0.2). We couldn’t estimate for other ethnic groups because of smaller sample sizes. Furthermore, the use of PCGC model which is designed for case/control approximately showed the same estimate (19.8%, SE=.07). However, the LDAK model completely showed different estimate (41.0%).

**Table 2.**
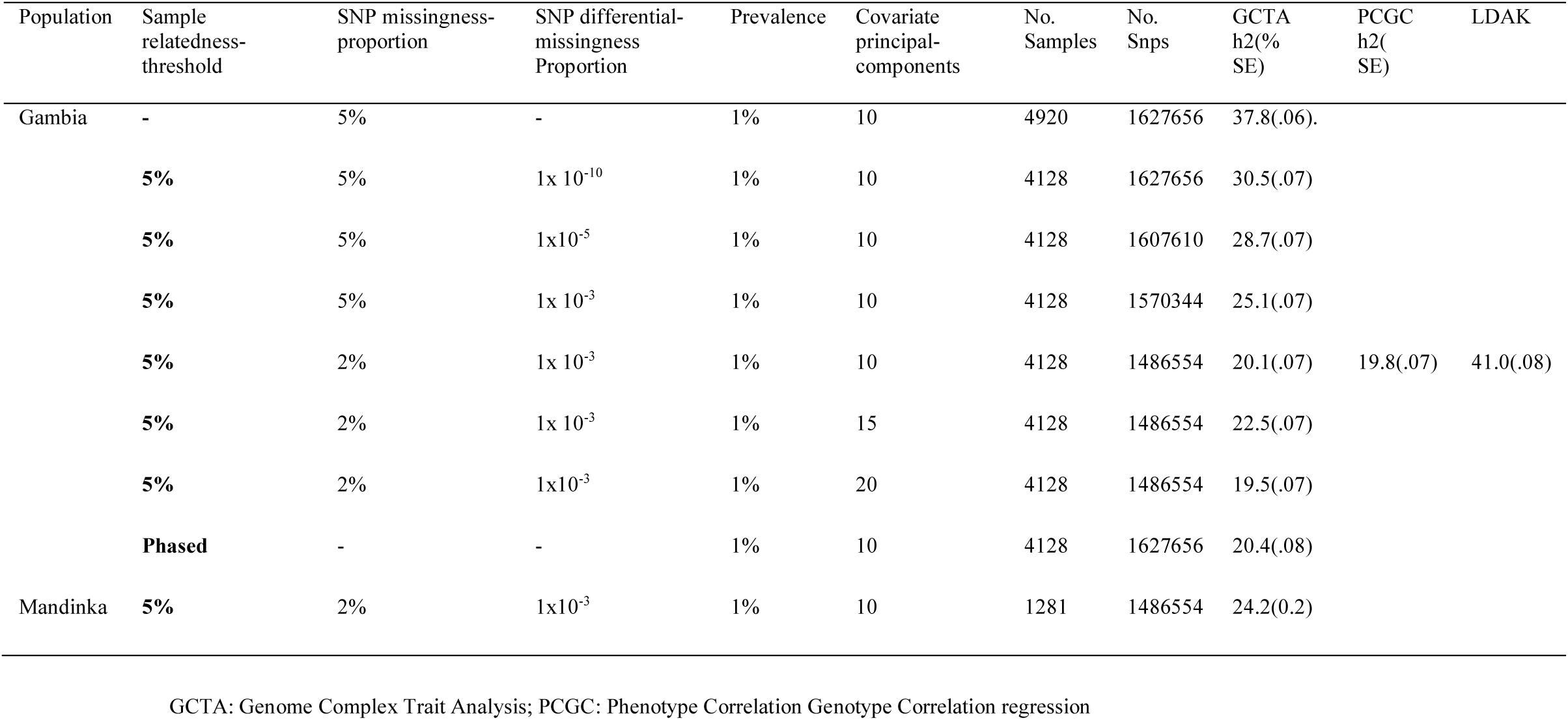
SNP-heritability of severe malaria susceptibility/resistance of Gambian population for different basic quality threshold using MLM

Encouraged by the fairly stable estimate, we partitioned the heritability of severe Malaria heritability in to intergenic and genic regions using GCTA model as shown in table (3). The proportion of heritability explained by a SNP in gene within 10kb boundary is on average twice as the heritability explained by a SNP in intergenic regions based on 10kb boundaries. Larger sample size is needed to further partition the heritability explained by cell types and functional groups.

**Table 3.** Partitioning of heritability in to genic and intergenic genomic regions based on10 kb boundaries using UCSC annotation data bases

|  | 10 Kb boundaries of genes |  |
| --- | --- | --- |
|  | Genic | Inter-genic |
| No. of SNPs | 827928 | 685896 |
| h<sup>2</sup>(%SE) | 16.3 (.06) | 6.4 (.05) |
| h<sup>2</sup> per SNP | 1.9x10<sup>-5</sup> | 9x10<sup>-6</sup> |

Although our heritability estimation is fairly stable when stringent QC is implemented, care should be taken in interpreting these results: First, all polygenic methods perform better in less structured data obtained from homogenous populations than the MalariaGen dataset which is comprised of diverse populations spanning most of the Malaria endemic belt in Africa. Second, the methods are designed and perform well in highly polygenic traits/diseases in which effects of each variant is mostly modest. However, a significant proportion of malaria protection trait might be attributed to rare variants of large effect sizes that might not be in LD with common variants and can’t not be ‘tagged’ by SNPs chip. The contribution of such variants will not be accounted for in the estimation. Third, subtle population structure that cannot be corrected by conventional methods such as unmatched case/control can potentially create systematic biases to the estimate. Fifth, the controversy between the performances of different methods is still continuing and there is no gold standard method against which the accuracy of these models can be evaluate

### Epistasis

Epistasis is becoming one of the hot research topics in genetic epidemiological studies in the last few years because of the fact that none additive genetic variations are shown to have significant influence on the phenotype of complex traits/diseases than previously expected^80^. The available statistical approaches and software packages for detection of epistasis at genome-wide scale are reviewed elsewhere^80,81^. These approaches have been applied in genetic studies of complex diseases such as lupus erythematosus^82^, anklosing spondlylitis^83^, psiorosis^84^ and unraveled previously unknown epistasis interactions between risk loci which explained a significant proportion of ‘missing’ heritability of the respective diseases.

Epistasis between malaria risk loci have been well documented and implicated as one of the possible reasons for lack of replication of susceptibility variants in different populations and the ‘missing’ heritability. For instance, sickle-cell trait (*HbS*) and *α thalassaemia* were shown to demonstrate negative epistatic interactions such that the protection against severe Malaria offered by *HbS* is reduced when co-inherited with *α+ thalassaemia*^85^.

A case-control study in Kenyan population also reported that *α+ thalassaemia* modulates the effects of Haptoglobin (Hp) variants in predicting the risk of severe malaria^86^. In this study, it was shown that the combination of *α*+ thalassaemia and Hp2-1 variant synergistically increase the protection against severe malaria by about 37%. However, the protective effect is decreased to 13% when *α*+thalassaemia is inherited with Hp1-1 and further diminished to neutral (zero) when inherited with *Hp2-2*. Similarly, in a mult-icenter case control study, the existence of negative epistasis interaction between *HbC* and *ATP2B* alleles was reported^^18^. Another recent case control study of severe malaria in Kenya reported the existence of negative epistasis between a compliment receptor called *S12* and *α+ thalassaemia* in which the protective effect of *S12* higher in children with normal *α-globin*87^. The extent and pattern of epistatic interaction at genome-wide scale is yet to explored. In malaria susceptibility GWAS, the priority has been given to a single locus analysis to identify novel risk loci. We expect that, the next step of Malaria susceptibility GWAS will be the considerations of epistasis.

## From GWAS to biology: multi**-**step and multi**-**dimensional analysis

### Fine-mapping and pathway analysis

The ultimate goal of genetic susceptibility studies is to identify the causative variants and understand the underlying biological pathways which can lead to translated medicine such as effective vaccines and therapeutics. However, translating GWAS signals in to biological themes remains an open problem because of the confounding effects from LD between association SNPs, limited knowledge of gene functions and localization of the majority of GWASs hits in none protein coding regulatory regions (regulatory SNPs)^^88–90^. In attempt to address this challenge, several fine mapping strategies have been developed and implemented91^. One such strategy is trans-ethnic fine mapping in which the natural variability of haplotype structure across ethnically diverse populations is used to narrow down the candidate causative variants^92^. The smaller LD and diversity of haplotype structure in African population makes it relatively easier to identify the causal SNPs and target gene/genes through fine mapping approaches^8^. However, the fact that malaria protective alleles are heterogeneous across populations might challenge the application of trans-ethnic fine-mapping approaches in malaria susceptibility studies.

Alternatively, several fine mapping statistical tools have recently been developed following the advances in annotation data bases and improved reference panels. These include Bayesian approach, heuristic approach and penalized regression methods ^93^. The principles, applications, strength and weakness of these methods is reviewed elsewhere ^91,93^. These methods are increasingly playing crucial role in the efforts being made to pinpoint causal variants of complex diseases/traits. For instance, Galarneau *et al*. ^94^identified novel independent association signals by fine-mapping three loci that are known to influence fetal hemoglobin (HbF) levels. The authors sequenced the three loci (*BCL11A, β- globin and HBS1L-MYB*), undertook dense genotyping, performed step-wise conditional analysis and revealed previously un-recognized SNPs that explain additional genetic variation. Similarly, a recent fine mapping study of HLA region identified several susceptibility loci for multiple infectious diseases^79^. More sophisticated studies that combine statistical and functional fine-mapping strategies have recently been implemented and provided mechanistic insights to the genetic basis of complex diseases^95^

In addition to fine mapping approaches, pathway and interaction analysis can be another avenue for exploring molecular basis of Malaria susceptibility/resistance. Instead of emphasizing on single-variant analysis, these approaches test the coordinated effects of several variants at systems level using biological information from annotation data basis ^96^. Pathway analysis improve study power by integrating cumulative effects of weak association signals and provide functional information by identifying associated sets of genes/proteins ^97^. By implementing the pathway analysis approaches, several studies have gained new insights in understanding the genetic basis of complex diseases^^98–100^.The available statistical tools for pathway and interaction analysis is reviewed in101^.We therefore, advocate for the implementation of fine mapping and pathway analysis in malaria susceptibility studies to shed more light to the underlying biology.

### Multi-omics approaches

Today, there are significant advances in high throughput technologies that can generate big ‘-omic’ data from all spectrum of molecular biology^88^. The ‘-omics’ studies (Genomics, Epigenomics, Transcriptomics, Proteomics, Metabolomics) are crucial to understand the underpinning biology of complex diseases.

In severe malaria, ‘-omics’ studies have provided important clues about the molecular events that lead to either complications or recovery from diseases. For instance, following the discovery of glycophorin regions by the GWAS, a targeted sequencing-based study ^31^was conducted to characterize variants in this region and identified a novel distinct copy number variant called *DUP4* which reduces the risk of severe Malaria by about 33%. In addition to this, a genome-wide gene expression study was conducted in Kenyan children in which increased expression of genes related to neutrophil activation during malaria infections was shown ^102^. The authors also observed differential expression of heme- and erythrocytes-related genes in acute malaria patients which reaffirms the importance of erythrocyte-related genes in malaria susceptibility/resistance.

In another host-parasite interaction study, the importance of miRNA in inhibiting parasite growth in erythrocyte was reported ^103^. The authors observed translocation of several host RBC miRNAs in to *P. falciparum* parasites, as well as fusion of these human miRNAs with parasite mRNA transcripts to inhibit the translation of enzymes that are vital for the parasite development. Specifically, two micro-RNAs, miRNA-451 and let-7i, were highly enriched in *HbAS* and *HbSS* erythrocytes and these miRNAS along with miR-223 were shown to attenuate the growth of parasite^103^.

However, ‘-omics’ studies are limited to single data-type analysis and lack adequate power to explain the complexity of molecular processes and usually lead to identification of correlations than causations^104^. Thus, integrating and analysing multiple ‘-omics’ data enables better understanding of the molecular processes and interactions that give rise to complex diseases/traits. For example, leveraging host microbiome relative abundance data as a second (quantitative) trait, and performing a joint analysis of bivariate phenotypes can increase statistical power by maximizing phenotypic information and inform how the interaction between host genotype with microbiome impacts the phenotype.

Multi-omics approaches aim to integrate big ‘-omics’ data, undertake ‘multi-step’ and ‘multidimensional’ analysis for elucidating complex biological problems^104^. Driven by the massive abundance of ‘-omics’ data from wide ranges of biological molecules, multi-omics strategy have recently provided unprecedented successes in complex diseases/trait studies. The current state of art of multi-omics approach and available statistical methods is recently reviewed in Hasin *et al.* ^104^

Malaria susceptibility/resistance is influenced by several host, parasite and environmental factors as depicted in Figure 1. The protective alleles have independently evolved in different populations being shaped by the co-evolution and interaction between the human genome, the parasite and environment^1^. So far, single ‘-omics’ data analysis enabled us to understand some of the factors that are associated with the malaria protective traits. To progress beyond associations and pinpoint the causal pathways, it may require to implement carefully designed, coordinated multi-omics studies that involve human host, the parasite, the environment and possibly mosquito. The current advents of high-throughput technologies in generating massive ‘-omics’ data and their continuously decreasing cost complemented with the availability of statistical tools which able to simultaneously capture millions of data points will lead to the implementations of multi-omics approaches in malaria susceptibility studies.

**Figure 1:**
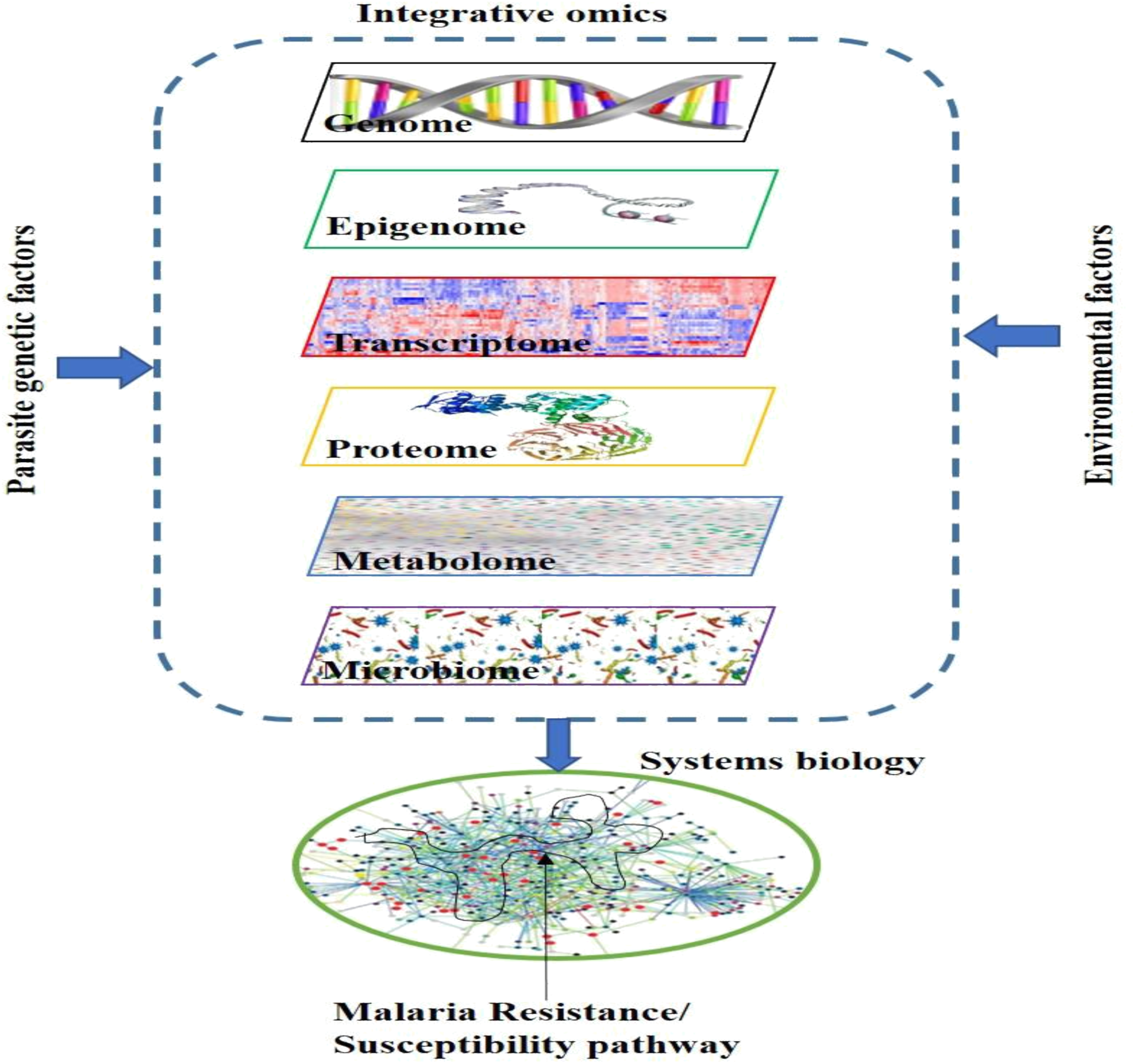
Schematic representation of the integrative analysis. Systems biology approach which incorporate multiple layers of information from host (multi-omics), environment and parasite genetic factors can potentially lead to the discovery of Malaria protective pathways

## Conclusions and perspectives

The ultimate goal of Malaria susceptibility study is to discover a novel causal biological pathway that provide protections against severe Malaria; a fundamental step towards translational medicine such as development of vaccine and new therapeutics that can facilitate the global malaria eradication efforts. To achieve this goal, several study approaches have been implemented at least for the last three decades and successfully identified association variants.

Recently, a number of GWASs have successfully been implemented in malaria endemic areas to better understand the underlying biology. While the well-known variants were replicated, only few novel variants were convincingly identified and their biological functions remains to be understood. Several limiting factors including genetic diversity of population in malaria endemic areas, allelic heterogeneity of protective variants, small sample sizes, lack of proper reference panel and proper genotyping chips might have negatively impacted the malaria GWASs.

Another challenge is that we don’t know much about the genetic architecture of malaria protective trait. There are two scenarios in which GWAS approach might fail. First, malaria protective trait might largely be attributed to rare variants of large effect sizes that might not be in LD with common variants and can’t not be ‘tagged’ by SNPs chip. Second, malaria might have left multiple genetic variants distributed across human genome; the majority of which have effects too small to be detected by the standard GWASs^1,7^. Theoretically, the large sample sizes, dense genotyping chips, use of appropriate reference panels and effective genotype imputation can address the majority of challenges. However, given the resource constraints; especially, in Africa where malaria problem is the greatest, this will likely take several years to achieve.

On the other hand, the recent advances in statistical techniques is enabling to extract useful information from the present-day GWAS sample sizes. For example, a number of statistical approaches have been developed to capture polygenicity in complex diseases. We showed how these methods can potentially be implemented in malaria susceptibility studies and provide useful insights. We believe that further studies with larger sample sizes can elucidate the polygenic effects in malaria protective trait by extending the analysis to genome partitioning, risk prediction and genetic correlations.

Beyond singe locus analysis, multi-step and multi-locus analysis including pathways analysis, fine mapping and interaction analysis can potentially be implemented in malaria susceptibility GWASs to gain new insights to the underpinning biology. For instance, pathway analysis can provide important information by analyzing the coordinated effects of several variants at systems level using biological information from annotation data basis. Fine mapping strategies that combine statistical and functional fine-mapping strategies can potentially be implemented to pinpoint the causal variants from the GWAS association signals.

Most importantly, the future direction of malaria susceptibility requires a paradigm shift from single ‘-omics’ to ‘multi-stage’ and ‘multi-dimensional’ integrative functional studies that combines multiple data types from the human host, the parasite, the mosquitoes and the environment. The current biotechnological advances, an ever-increasing annotation data bases and availability of advanced analytical techniques, will eventually lead to feasibility of systems biology studies and revolutionize malaria research.

## Acknowledgements

We thank Kwiatkowski’s group from University of Oxford for their constructive comments and assistance. We thank Kirk Rokett and Ellen Leffler for their useful comments. We are very grateful to Gavin Band for his supervision and guidance in heritability analysis and his critical comments. I thank Newton’s fund student transfer scheme for funding me during my stay at University of Oxford. We also thank Abdoulaye Djimde for his follow up.

## Authors contributions

DD designed and drafted the manuscript; LG and AD revised the manuscript; EC revised the manuscript and supervised the work

